# Heat stress modifies the lactational performances and the urinary metabolomic profile related to gastrointestinal microbiota of dairy goats

**DOI:** 10.1101/385930

**Authors:** Alexandra Contreras-Jodar, Nazri Nayan, Soufiane Hamzaoui, Gerardo Caja, Ahmed A.K. Salama

## Abstract

The aim of the study was to identify the candidate biomarkers of heat stress (HS) in the urine of lactating dairy goats through the application of proton Nuclear Magnetic Resonance (^1^H NMR)-based metabolomic analysis. Dairy does (n = 16) in mid-lactation were submitted to thermal neutral (TN; indoors; 15 to 20°C; 40 to 45% humidity) or HS (climatic chamber; 37°C day, 30°C night; 40% humidity) conditions according to a crossover design (2 periods of 21 days). Thermophysiological traits and lactational performances were recorded and milk composition analyzed during each period. Urine samples were collected at day 15 of each period for ^1^H NMR spectroscopy analysis. Principal component analysis (PCA) and partial least square–discriminant analysis (PLS-DA) assessment with cross validation were used to identify the goat urinary metabolome from Human Metabolome Data Base. HS increased rectal temperature (1.2°C), respiratory rate (3.5-fold) and water intake (74%), but decreased feed intake (35%) and body weight (5%) of the lactating does. No differences were detected in milk yield, but HS decreased the milk contents of fat (9%), protein (16%) and lactose (5%). Metabolomics allowed separating TN and HS urinary clusters by PLS-DA. Most discriminating metabolites were hippurate and other phenylalanine (Phe) derivative compounds, which increased in HS vs. TN does. The greater excretion of these gut-derived toxic compounds indicated that HS induced a harmful gastrointestinal microbiota overgrowth, which should have sequestrated aromatic amino acids for their metabolism and decreased the synthesis of neurotransmitters and thyroid hormones, with negative impact on milk yield and composition. In conclusion, HS markedly changed the thermophysiological traits and lactational performances of dairy goats, which were translated into their urinary metabolomic profile through the presence of gut-derived toxic compounds. Hippurate and other Phe-derivative compounds are suggested as urinary biomarkers to detect heat stressed dairy animals in practice.

## Introduction

Exposure to high ambient temperature induces several physiological responses in order to maintain body homeostasis. Animals suffer from heat stress (HS) when physiological mechanisms fail to counterbalance an excessive heat load [1]. Exposure of dairy animals to HS results in a decline in their productive [2] and reproductive [3] performances due to a strong metabolic disruption. Dairy animals under HS typically show decreased feed intake, increased water consumption and thermophysiological traits, such as respiratory rate and rectal temperature, when compared to thermoneutral (TN). Usually, HS reduces milk yield and impairs milk composition in dairy goats [4]. Although these negative effects on milk production are traditionally attributed to a decline in feed intake, pair-fed TN experiments have shown that intake only accounts for 35-50% of milk yield reduction in dairy cows [5, 6]. Therefore, there is a specific effect of HS that disrupts body metabolism and milk secretion which remains unknown.

Biofluid assessment by Nuclear Magnetic Resonance (NMR) spectroscopy can shed some light on the physiological mechanisms occurred in animals when exposed to HS. Proton (^1^H) NMR, together with multivariate statistical analysis, has been successfully used as metabolite profiling method to study the metabolic changes in HS rats [7]. This robust and reliable technique provides vast information on metabolome dynamics and metabolic pathways [8]. The ^1^H NMR spectra derives from thousands of metabolite signals that usually overlap, adding complexity to data processing. Computer-based data reduction and multivariate statistical pattern recognition methods, such as principal component analysis (PCA) and partial least square–discriminant analysis (PLS-DA), have shown to be beneficial techniques to get the most from the information obtained in the ^1^H NMR spectra for classification purposes [9, 10].

To our knowledge, no studies have been carried out to evaluate urine metabolomics of dairy goats. The aim of this study was to identify the candidate biomarkers of HS through the application of ^1^H NMR-based metabolomic urinalysis of dairy goats.

## Material and methods

### Animals and treatments

Animal care conditions and management practices agreed with the procedures stated by the Ethical Committee of Animal and Human Experimentation (reference CEEAH#09/771) of the Universitat Autonoma of Barcelona (UAB) and the codes of recommendations for livestock wellbeing of the Ministry of Agriculture, Food and Environment of Spain.

Sixteen multiparous Murciano-Granadina dairy does (43.5 ± 1.6 kg body weight), lactating and open, from the herd of the SGCE (Servei de Granges i Camps Experimentals) of the UAB in Bellaterra (Barcelona, Spain), were blocked in 2 balanced groups at mid-lactation (81 ± 3 days-in-milk; 2.00 ± 0.04 L/day). Does were adapted to metabolic cages for 2 weeks before the start of the experiment and the groups randomly allocated to 2 ambient conditions treatments according to a 2 × 2 (treatment × period) crossover design. Treatments were TN (indoors shelter; 15 to 20°C and 45 ± 5% relative humidity) and HS (climatic chamber 4 × 6 × 2.3 m with temperature-humidity control system, Carel Controls Ibérica, Barcelona, Spain; 37 ± 0.5°C during the day, and 30 ± 0.5°C during the night; 40 ± 5% humidity and 90 m^3^/h continuous air turnover). Day-night length was set to 12-12 hours and the experimental periods lasted 21 days (14-days adaptation, 5-days measurements, 2-days washout). Temperature-humidity index (THI) calculated according to NRC (1971) resulted THI_TN_ = 59 to 65 and THI_HS_ = 75 to 83. Experimental conditions were similar to those detailed by Hamzaoui et al. [11].

Does were milked once-a-day (0800) with a portable machine (Westfalia-Separator Ibérica, Granollers, Spain) set at 42 kPa, 90 pulses/min and 66% pulsation ratio and provided with volumetric recording jars (3 L ± 5%). The milking routine included cluster attachment, machine milking, machine stripping before cluster removal, and teat dipping in an iodine solution (P3-ioshield, Ecolab Hispano-Portuguesa, Barcelona, Spain). Feed was offered ad libitum at 0930 (130% feed intake of the previous day) and consisted of a total mixed ration (dry matter, 89.9%; net energy for lactation, 1.40 Mcal/kg; crude protein, 17.5%; organic matter, 87.3%; neutral detergent fiber, 34.4%; acid detergent fiber, 21.8%; on dry matter basis). Ration ingredients were (as fed): alfalfa hay, 64.2%; ground barley, 9.6%; beet pulp, 9.6%; ground corn, 8%; soybean meal, 3.3%; sunflower meal, 3.2%; molasses, 1%; salt, 0.6%; sodium bicarbonate, 0.3%; mineral and vitamin complex, 0.2% (Vitafac premix, DSM Nutritional Products, Madrid, Spain). Water was permanently available and offered at room temperature in water bowls connected to individual tanks of 20 L. A sawdust drip tray under each water bowl was used to collect spilled water.

## Sampling and measurements

### Thermophysiological traits and lactational performances of the goats

Does were weighed at the start and the end of each period using an electronic scale (True-Test SR2000, Pakuranga, New Zealand; accuracy, 0.2 kg). Rectal temperature (digital clinical thermometer, ICO Technology, Barcelona, Spain; accuracy, 0.1°C) and respiratory rate (flank movements during 60 s) were recorded daily at 0800, 1200, and 1700 throughout the experiment. Milk yield (volume) was recorded daily throughout the experiment and milk samples collected weekly for composition (NIRSystems 5000, Foss, Hillerød, Denmark). Feed and water intakes were calculated by weight from the daily refusals and feed samples were collected daily and composited by period for analyses. Feed composition was determined according to analytical standard methods [12].

### Urine sampling and preparation

Urine samples from each doe were collected at micturition on the morning of day 15 of each period (n = 32) and stored at −20°C until ^1^H NMR analysis.

Preparation of samples for ^1^H NMR spectroscopy was done according to Beckonert et al. [8]. Briefly, phosphate buffer solution (pH 7.4) was prepared with sodium phosphate dibasic (Na_2_HPO_4_; 99.95% trace metals basis, anhydrous, Sigma-Aldrich, Germany), sodium phosphate monobasic (NaH_2_PO_4_; 99.95% trace metals basis, anhydrous, Sigma-Aldrich) and sodium azide (NaN_3_; Sigma-Aldrich). Deuterium oxide (D_2_O; 99.9 atom % D, Sigma-Aldrich), containing 0.75% 3-(trimethylsilyl) propionic-2,2,3,3-d4 acid (TSP) sodium salt as NMR reference compound, was added before the flask was filled up to 25 mL with milli-Q water (EMD Millipore, Darmstadt, Germany). The flask was shaken thoroughly and left in a Clifton sonicator (Nickel Electro, Weston-super-Mare, United Kingdom) at 40°C until the salts were dissolved. The prepared phosphate buffer solution was stored at 4°C. Urine samples were thawed in a water bath, thoroughly shaken and spun for 5 min at 12,000 × *g* in a swing-bucket rotor (Hettich, Tuttlingen, Germany) at 4°C. Then, 400 μL of the urine sample were transferred into Eppendorf tubes and mixed thoroughly with 200 μL of cold phosphate buffer solution. All the tubes were then centrifuged for 5 min at 12,000 × *g* at 4°C and 550 μL of the final mixture transferred into 5-mm NMR tubes (VWR International Eurolab, Barcelona, Spain). The prepared NMR tubes were immediately put on ice and sent to the NMR Service of the UAB for ^1^H High Resolution NMR Spectroscopy.

### NMR spectroscopy

^1^H NMR spectra were acquired on a Bruker Avance-III spectrometer (Bruker BioSpin, Rheinstetten, Germany) operating at a ^1^H NMR frequency of 600 MHz and a temperature of 298°K, controlled by a Burner Control Unit-extreme regulator. A 5 mm Triple Resonance Broadband Inverse probe with *z*-gradients and inverse detection was used and controlled by TopSpin2.1 software (Bruker, Germany). One-dimensional ^1^H NMR spectra were obtained using an one-dimensional Nuclear Overhauser Enhancement Spectroscopy (NOESY) pulse sequence. The solvent signal was suppressed by pre-saturation during relaxation and mixing time. A total of 32 scans and 2 dummy scans were performed to produce 32,768 data points for each spectrum using a relaxation delay of 2.0 s with a pulse power level of 54 dB and an acquisition time of 2.6 s. Spectral width (δ) used for all data collected was 12.0 ppm, and 0.3 Hz exponential line broadening was applied for the Fourier-transform of the raw data. ^1^H NMR spectra were phased, baseline corrected, and corrected for chemical shift registration relative to the TSP reference compound previously indicated (δ = 0.0 ppm) in TopSpin 2.1.(data in S1 File).

## Statistical analyses

### Thermophysiological and performance analysis

Data were analyzed by the PROC MIXED for repeated measurements of SAS v. 9.1.3 (SAS Inst. Inc., Cary, North Carolina, USA). The statistical mixed model contained the fixed effects of environmental treatment (TN vs. HS), the period (1 and 2) and measuring day (1 to 19), and the random effects of the animal (1 to 16), the interactions (treatment × day and treatment × period), and the residual error. Differences between least squares means were determined with the PDIFF option of SAS. Significance was declared as P<0.05.

### NMR data pre-processing and analysis

Pre-treatment of raw spectral data is critical for generating reliable and interpretable models using multivariate analysis techniques. Nevertheless, metabolic fingerprinting datasets acquired from ^1^H NMR spectrometers suffer from imprecisions in chemical shifts due to temperature, pH, ionic strength and other factors. Therefore, models generated using multivariate analysis may fail to identify separations between classes, and their loadings can be difficult to interpret due to over-abundance of variables. To mitigate these complications each spectrum was uniformly divided into ‘bins’ and the signal intensities within each bin were integrated to produce a smaller set of variables according to Worley and Powers [13]. In this way, each dataset was divided in bins of 100 (i.e., from 0.0003 to 0.03 ppm) using R software v. 3.2.3 [14]. After binning, alignment and normalization of spectra were accomplished to ensure that all observations were directly comparable. In this sense, urine spectra were normalized to creatinine methyl resonance intensity at δ = 3.05 ppm and then log_2_ transformed. Regarding variable selection, raw ^1^H NMR spectral data were edited by excluding the regions outside the chemical shift range of δ *=* 9.0-0.5 ppm, and also the residual peak of the imperfect water suppression (δ = 5.5-4.6 ppm). Following the recommendations of Pechlivanis et al. [15], the spectral regions of histidine, 1-metylhistidine, and 3-methylhistidine (δ = 8.17-7.87, δ = 7.15-7.01, and δ = 3.77-3.71 ppm, respectively) were also removed because of the sensitivity to small pH differences among urine samples.

Once ^1^H NMR pre-processing data was completed, data were subjected to multivariate statistical analysis. Initially, PCA was performed without considering the class information for samples examination and search for outliers. Then, PLS-DA with leave-one-out cross validation was also performed on the datasets using the pls package of R software [16]. PLS-DA allowed individual samples to be classified according to the respective class prior to analysis (TN or HS). Model strength was assessed using both R^2^ and Q^2^ statistical parameters. While R^2^ values reported the total amount of variance explained by the model, the Q^2^ reported model accuracy as a result of cross-validation. Aside from its theoretical maximum value of 1, for biological models, an empirically inferred acceptable value is ≥ 0.4 [9]. The resulting Q^2^ statistic was compared to a null distribution to test model significance (P<0.05).

Interpretation of multivariate analysis was performed through scores and loadings plots according to its contribution to the separation between groups. For biomarker search, PLS-DA loadings were sorted by absolute values, being the first ones the metabolites responsible of the separation between experimental groups. Because both PCA and PLS-DA analysis may be influenced by variable correlations, the intra- and the inter-class variance of metabolites may have no significant differences in the one-dimensional statistical analysis [17]. For this reason, spectral bins were also selected using a Volcano plot with paired student *t* test analysis between HS over TN cohorts to get a general overview of the data (log_2_ fold change thresholds, ≤1.5 and ≥1.5; P<0.01). All ^1^H NMR data pre-processing, statistical analysis and the generated plots were performed using R software v. 3.2.3 [14].

### Metabolite assignment

Chemical shifts linked to the highest loading values found in PLS-DA were annotated for metabolite assignment as HS biomarker candidates. The candidate chemical shifts and corresponding metabolites were assigned using the Human Metabolome Database [18] and queried in KEGG (Kyoto Encyclopedia of Genes and Genomes) database to know in which metabolic pathways they were involved.

## Results and discussion

### Effects of heat stress on thermophysiological and lactational performances of the goats

The effects of the experimental HS conditions on thermophysiological and lactational performances of the dairy goats are summarized in Table 1. Rectal temperature and respiratory rate increased during the day in both groups of does, following the expected circadian rhythm and the daily THI pattern in both TN and HS conditions. The greatest values were observed in the HS does at 1700, the increases being 1.2°C and 3.5-fold (P<0.001) when compared to TN does. On average, feed intake decreased 35% in HS (P<0.001) when compared to TN does but, in contrast, water consumption increased 74% (P<0.001). Furthermore, HS does lost 115 g/d of body weight, whereas TN goats won 162 g/d, on average (P<0.001). Obtained results agreed with those reported for the same breed of dairy goats in late-lactation and under similar HS conditions [11].

**Table 1.** Thermophysiological and lactational performances of dairy goats under thermal neutral (TN) and heat stress (HS) conditions. Values are least square means and standard error of the means (SEM).

| Item | Treatment |  | SEM | P value |
| --- | --- | --- | --- | --- |
|  | TN | HS |  |  |
| **Rectal temperature, °C** |  |  |  |  |
| 0800 | 38.5 | 39.1 | 0.08 | <0.001 |
| 1200 | 38.7 | 39.7 | 0.07 | <0.001 |
| 1700 | 38.7 | 39.9 | 0.09 | <0.001 |
| **Respiratory rate, breaths/min** |  |  |  |  |
| 0800 | 27 | 69 | 4 | <0.001 |
| 1200 | 39 | 131 | 6 | <0.001 |
| 1700 | 37 | 130 | 6 | <0.001 |
| **Performances** |  |  |  |  |
| Dry matter intake, kg/d | 2.26 | 1.47 | 0.09 | <0.001 |
| Water intake, L/d | 6.1 | 10.6 | 1.0 | <0.001 |
| Final body weight, kg | 48.6 | 39.8 | 1.8 | <0.001 |
| Body weight variation, kg | 3.5 | -2.1 | 1.0 | <0.001 |
| Milk yield, L/d | 1.88 | 1.79 | 0.11 | 0.413 |
| FCM<sup>1</sup> yield, L/d | 2.17 | 1.86 | 0.13 | 0.017 |
| **Milk composition, %** |  |  |  |  |
| Fat | 3.98 | 3.64 | 0.13 | 0.009 |
| Protein | 3.40 | 2.85 | 0.10 | <0.001 |
| Lactose | 4.51 | 4.30 | 0.07 | 0.003 |

<sup>1</sup>Fat corrected milk at 3.5%; $FCM = L \times [0.432 + 0.162 \times (\text{fat, \%})]$ , being L liters of milk.

Reducing feed intake is a way to decrease heat production in warm environments because heat increment of feeding, especially in ruminants, is an important source of heat production [4]. Moreover, increased water consumption under HS conditions is mainly used for boosting latent heat losses by evaporation (e.g., sweating and panting). Despite this, no differences in milk yield were observed, although milk composition markedly worsened. Milk fat, protein and lactose contents varied by −9%, −16% and −5%, respectively (Table 1; P<0.01), which would severely compromise the milk transformation to dairy products [19]. Consequently with the decrease in milk composition, fat corrected milk yield also varied by −14% (P<0.05).

Although our does were less sensitive to HS than dairy cows with regard to feed intake and milk yield, the effects of HS on milk fat content and fat corrected milk were contradictory when compared to cows. So, despite the typical fat depression seen in commercial dairy cow farms during the summer, Rhoads et al. [5] and Shwartz et al. [20] reported 9% increase or no change in milk fat content, at the short- or mid-term, respectively, in HS vs. TN dairy cows. On the other hand, the above indicated negative effect of HS on the milk protein content of our goats (i.e., −16%), was greater than reported by Rhoads et al. ([5], −5%) and Shwartz et al. ([20], −9%) in dairy cows, and herewith in the same breed of dairy goats in late lactation ([11], −13%). The negative effects of HS on the lactational performances of dairy ruminants are usually attributed to the decline in feed intake, but pair-fed experiments under TN conditions have shown that feed intake only explains approximately a half of the fall in milk yield and body weight in dairy cows [5, 6]. Therefore, similar responses were expected in our dairy goats.

As an intermediate conclusion, the thermophysiological and lactational performance responses observed, clearly evidenced that our HS does (kept at THI = 75 to 83) were under severe stress on the days at which the urine samples for ^1^H NMR-metabolomics assessment were collected (day 15).

### NMR urinary spectroscopy of the goats

Comparison of ^1^H NMR urinary mean spectra for the TN and HS lactating does is shown in Fig 1. Resonance assignments reported in the figure were made from the known chemical shifts and coupling patterns of urine spectra previously described in humans [14, 21].

**Fig 1.**
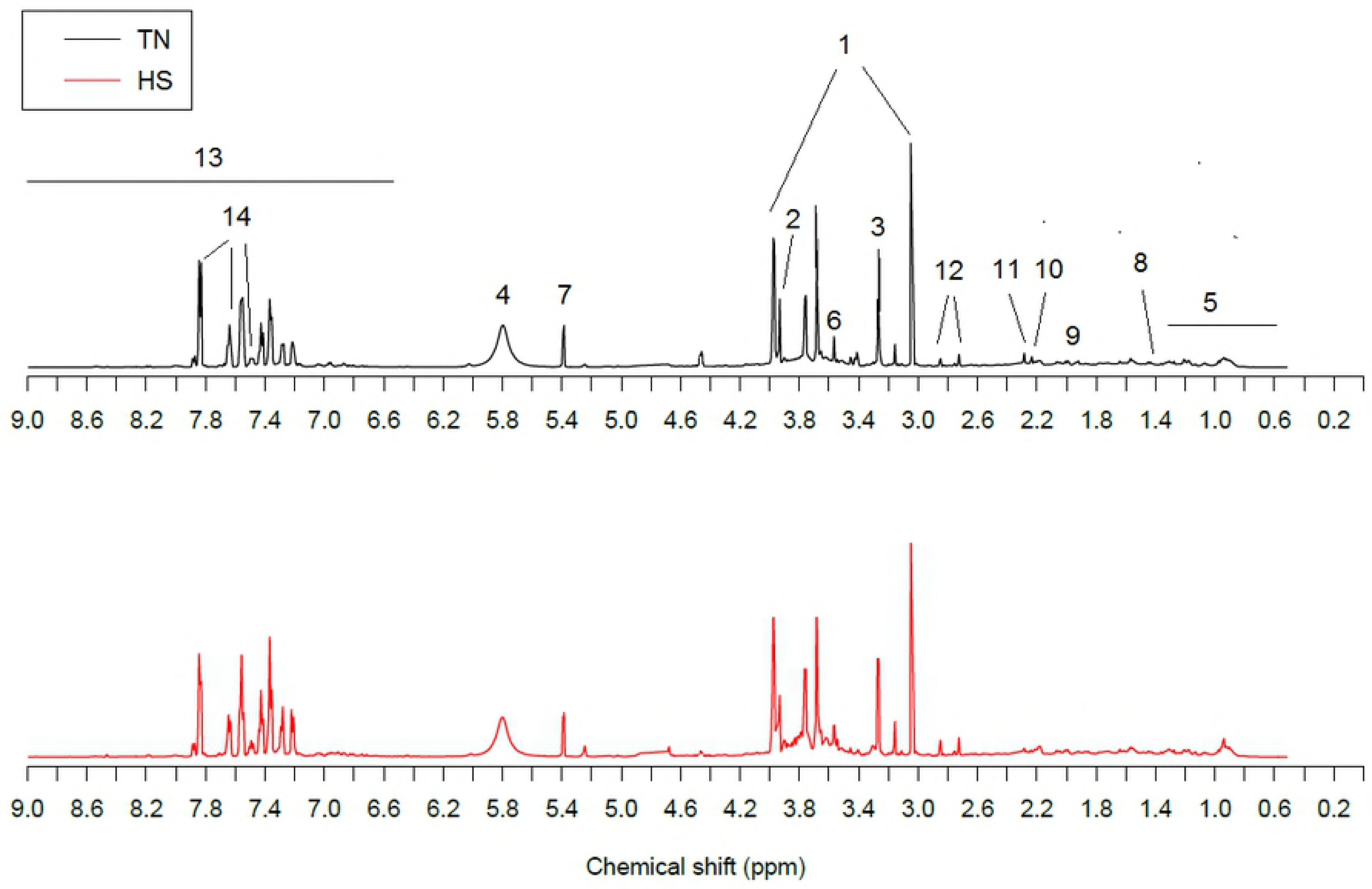
One-dimensional ^1^H NMR spectra at 600 MHz of urine from representative thermoneutral (TN) and heat stressed (HS) dairy does. Dominant metabolites were: 1, creatinine; 2, creatine; 3, trimethyl-N-oxide; 4, urea; 5, branched-chain amino acids and organic acids; 6, glycine; 7, allantoin; 8, alanine; 9, N-acetyl glycoprotein; 10, glutamate; 11, succinic acid; 12, citric acid; 13, aromatic signals; 14, hippuric acid.

At first sight, visible differences in urine metabolites were found between HS and TN groups. Spectral region from δ = 8.0-6.5 ppm showed higher excretion compounds in the HS doe group. On the contrary, all excreted compounds that lay on the δ = 4.5-0.5 ppm spectral region appeared to be at lower concentrations in the HS group. More detailed analysis of metabolic differences between these two thermal conditions were obtained from the multivariate PCA and PLS-DA data analyses and the Volcano plot.

First, the Volcano plot (Fig 2) showed that TN does excreted a greater number of urinary metabolites (i.e., higher number of left-sided spots) than HS does. Most probably, this was a consequence of the metabolic sparing of nutrients of the HS does, which loosed weight as a result of their negative energy balance, to cope with the HS conditions.

**Fig 2.**
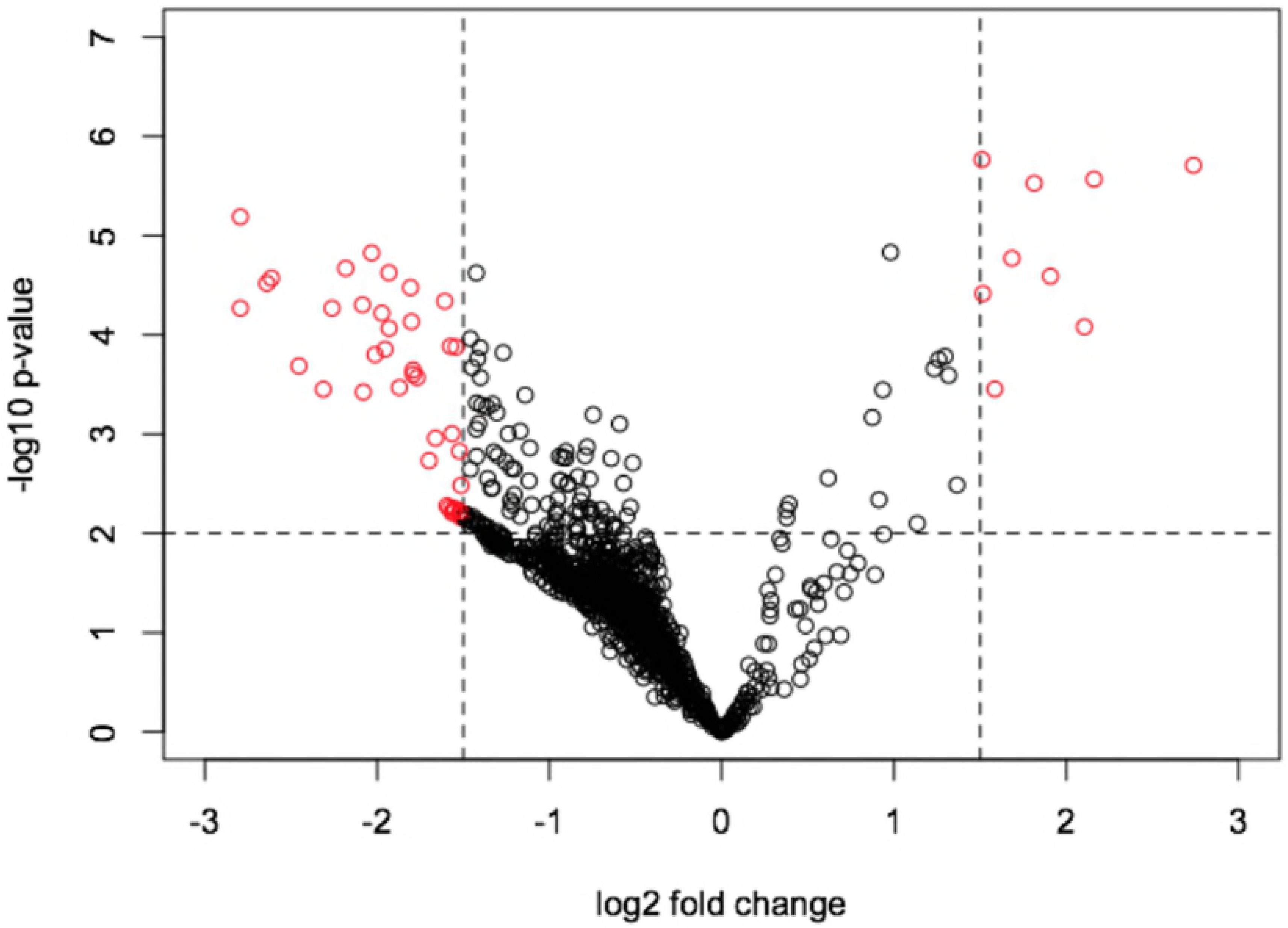
Volcano plot based on fold change (log_2_) and P value (−log_10_) of all spectral bins of ^1^H NMR urinary spectroscopy of thermoneutral (TN) and heat stressed (HS) lactating dairy does. Red circles indicate the spectral bins that showed significant changes and absolute fold changes greater than 1.5.

Regarding the multivariate analysis, PCA was initially applied to the ^1^H NMR spectra. Based on the principle of minimum differentiation, no samples were identified as outliers according to Hotelling’s T^2^ (95% interval of confidence). Therefore, all samples remain for subsequent PLS-DA in order to identify the metabolic differences between HS and TN dairy does. The PLS-DA scores plot showed a slight distinguishable separation between HS and TN datasets (Fig 3).

**Fig 3.**
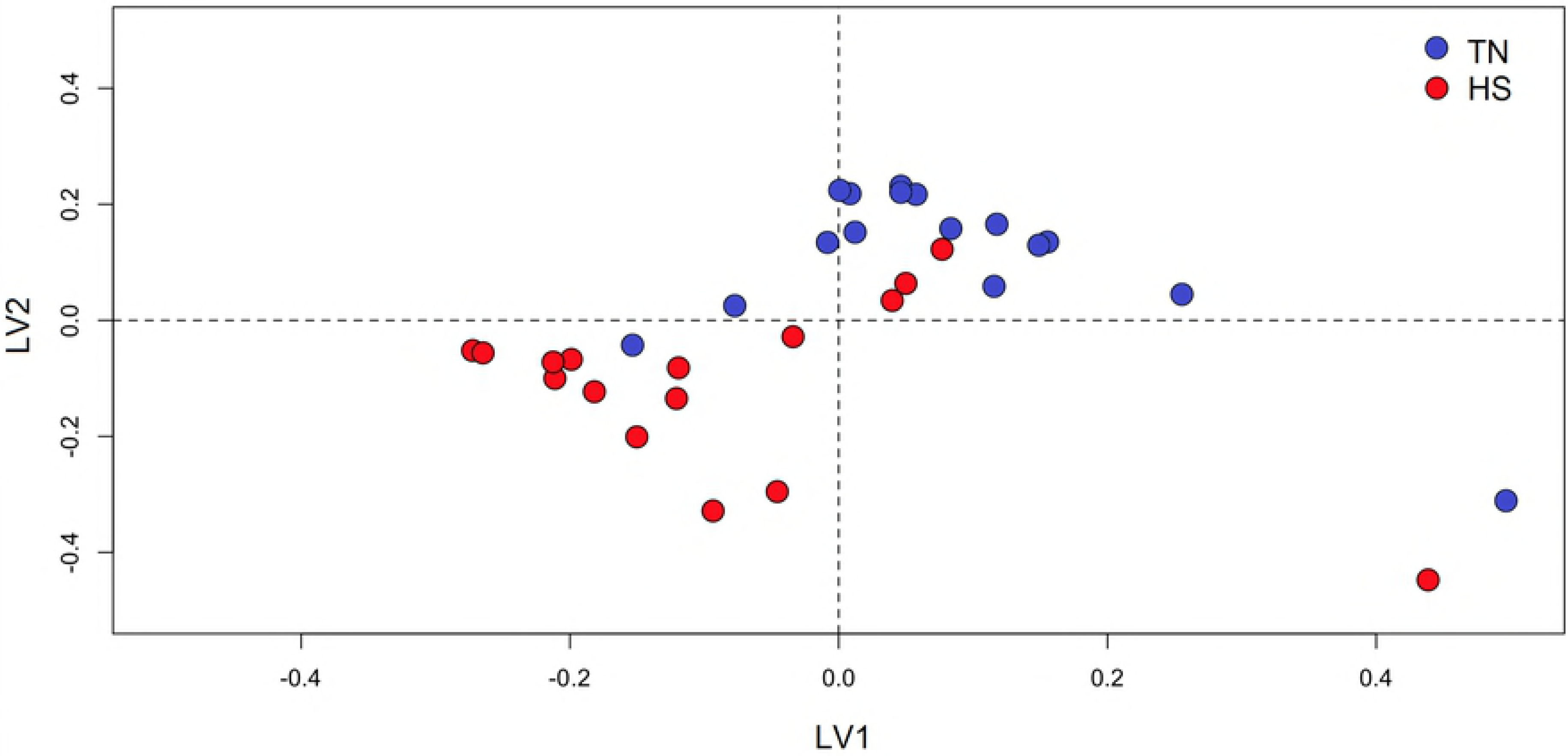
PLS-DA scores plot of the first two principal components of ^1^H NMR urinary spectra of thermoneutral (TN) and heat stressed (HS) lactating dairy does.

The separation along the x-axis (PLS-DA component 1) represents differences related to the environmental treatment. All other variations in the NMR data are visualized as separation in the y-axis direction (second component). The cross-validation of plasma metabolomics PLS-DA models (first 2 components) gave R_x_^2^ = 0.54, R_y_^2^ = 0.17, and Q^2^ = 0.47. The R^2^ and Q^2^ values in the model were higher than in the random model (P<0.01). The top-ranking urinary metabolites responsible for discriminating HS does were hippurate (δ = 7.54 ppm, 7.83 ppm and 7.63 ppm) and the phenylalanine (Phe) derivative compounds OH-phenylacetate (δ = 7.27 ppm), OH-phenylacetylglycine (δ = 7.20 ppm) and phenylglyoxylate (δ = 7.62 ppm). All of them were excreted in greater concentrations in the urine of HS when compared to TN does. Table 2 shows the excreted metabolites with the greatest quantitatively change under HS conditions, according to the results of the univariate and multivariate analyses.

**Table 2.** Selected metabolites contributing to the classification of the urine metabolome of thermoneutral and heat stressed lactating dairy does.

| Pathway | Chemical shift ($\delta$, ppm) | Metabolite | Fold change* | P value |
| --- | --- | --- | --- | --- |
| Phenylalanine (Phe) metabolism | 7.83, 7.63, 7.54 | Hippurate | 2.74 | <0.001 |
| Microbial metabolism in diverse environments | 7.62 | Phenylglyoxylate | 1.59 | <0.001 |
|  | 7.27 | OH-phenylacetate | 1.51 | <0.001 |
| Tyrosine (Tyr) metabolism | 7.20 | OH-phenylacetyl glycine | 2.16 | <0.001 |

\* Metabolites with positive fold change values means that they are excreted in greater concentrations under heat stress conditions

As it can be observed, of them come from certain gastrointestinal bacteria that generate uremic toxins that are absorbed into the blood and mainly cleared by the kidney [22, 23]. The greater excretion of these metabolites, also known as gut-derived uremic toxins and mammalian-microbial cometabolites [24], may be related to an abnormal overgrowth of gastrointestinal microbiota that is commonly accompanied with intestinal hyper-permeability [23, 25, 26].

The increase of gut-derived uremic toxins reflected alterations in the gastrointestinal environment due to the metabolic impact of HS. In fact, it is well known that under HS conditions, mammals redistribute blood to the periphery for heat dissipation purposes, while vasoconstriction occurs in the gastrointestinal tract [27] that leads to tissue hypoxia and oxidative stress [28]. Moreover, lower rumen pH has been reported as a side effect in HS goats [29] and it may compromise the integrity of the gastrointestinal tract barrier [2].

Hippurate and other Phe-derivative compounds are produced by the aerobic and anaerobic degradations of aromatic amino acids (e.g., Phe and Tyr) and dietary polyphenols by the gastrointestinal microbiota [22, 30, 31]. Although they are usually found in plasma and excreted in urine by crossing the cellular and tissue barriers (gastrointestinal epithelium, lymphatic barrier and liver), high levels of Phe-derivatives in urine are related to gastrointestinal leaky stages [32]. Moreover, high levels of gut-derived uremic toxins seem to affect both the cellular protein expression and the activity of the cyclooxygenase-2 (COX-2) enzyme, which plays a major role in the regulation of inflammation trough the production of prostaglandins; so, when COX-2 activity is speed up, inflammation increased [33]. Some Phe-derivatives also produce cytotoxic effects by the inhibition of cell pores opening and the production of reactive oxygen species [34].

Among Phe-derivatives, hippurate has a strong association with diet and gastrointestinal microbiota and its production requires of both microbial and mammalian metabolisms [24]. Gastrointestinal bacteria produce benzoic acid from dietary aromatic compounds, which is absorbed into the blood. Because of the toxicity of benzoic acid, it is conjugated with glycine in the mitochondrial matrix of the liver and renal cortex to form hippuric acid [24, 25], which is later filtered in the kidneys and finally excreted in urine as hippurate [24, 35]. The main elimination route for hippurate is the active renal tubular secretion and its disruption results in its accumulation in the blood [24]. Hippurate is a uremic toxin that participates in the correction of metabolic acidosis by stimulating ammoniagenesis, a dominant and adaptive mechanism of proton excretion. Moreover, it interferes several metabolic processes, such as: inhibition of glucose utilization by the kidney and muscle, modulation of fatty acid metabolism and regulation of acid-base balance by stimulating kidney’s ammoniagenesis, among others, as reviewed by Dzúrik et al. [35].

It might also be noted that, additionally to the production of gut-derived uremic toxins from dietary aromatic amino acids by the gastrointestinal microbiota, Phe is known to be an essential amino acid for most animals, including ruminants [36]. It is also the precursor of Tyr, that is essential for the synthesis of thyroid hormones and levodopa neurotransmitter. Previous studies have evidenced a strong decrease in plasma thyroid hormones (i.e., TSH, T4 and T3) of steers under HS conditions [37], which means that the basal heat production may in fact decrease when Phe and Tyr are scarce. Moreover, the rate of milk production is markedly affected by thyroid hormones, which modulate the nutrient partitioning towards milk production [38]. On the other hand, a decrease in the dopaminergic neurons activity was also observed in HS calves [39]. The drop of levodopa synthesis may be the result of the hypersecretion of its antagonist prolactin, as observed in response to HS in goats [40], ewes [41] and cows [42]. Prolactin is not only a hormone related to milk production, but has a broad variety of biological functions related to thermoregulation and water balance. The increase in plasma prolactin is not reflected in an increase in milk yield, as seen in dairy ruminants under HS conditions [11, 43]. Alamer [44] concluded that the mammary gland experiences a down-regulation of prolactin signaling pathways that could partially explain the depressed milk production of dairy cows during HS.

## Conclusions

Heat stress caused marked changes in thermophysiological traits and lactational performances of dairy goats, which were translated into their ^1^H NMR metabolomic urinary profile. These changes were related to the over-excretion of gut-derived toxic compounds generated by the gastrointestinal microbiota with expected decreases in the bioavailability of aromatic amino acids and impairment of the synthesis of thyroid hormones and neurotransmitters (i.e., levodopa, serotonin), which compromised the milk production of dairy goats. In practice, the use of hippurate and other phenylalanine derivatives are suggested as urinary biomarkers to identify heat stressed animals.

## Acknowledgements

The authors are grateful to the technical team of SGCE (Servei de Granges i Camps Experimentals) of the UAB for the care of the animals.

## Supporting information

S1 Dataset. ^1^H NMR data matrix of normalized and binned spectral data. HS, heat-stressed lactating dairy does; TN, thermal neutral lactating dairy does.

S1 Table. ^1^H NMR data matrix of normalized and binned spectral data. HS, heat-stressed lactating dairy does; TN, thermal neutral lactating dairy does.

